# Neutrophil progenitor cell therapy rescues host defense against *Staphylococcus aureus* in murine chronic granulomatous disease

**DOI:** 10.1101/2025.03.26.645550

**Authors:** Kristina D. Hinman, Jason T. Machan, Craig T. Lefort

## Abstract

Despite advances in engineered adaptive immune cell therapies, current options for innate immune cell therapies are sparse. In this work, we demonstrate the utility of a neutrophil progenitor-based cell therapy. Murine conditionally-immortalized neutrophil progenitors (NPs) overcome the hurdles of alternative cell therapies by engrafting in the unconditioned host and substantially contributing to the host neutrophil population. Here we demonstrate the therapeutic value of NPs using a murine model of the primary immunodeficiency chronic granulomatous disease (CGD). Those with CGD are highly susceptible to infection with *Staphylococcus aureus* because of genetic mutations that impair neutrophil antimicrobial function. We find that the prophylactic treatment of CGD mice with transfused NPs rescue them from an otherwise lethal *S. aureus* pulmonary infection. In investigating the mechanisms behind the improved clearance of *S. aureus* and survival of CGD mice, our data suggests that the antimicrobial function of host CGD neutrophils is rescued by the presence of donor-derived wild-type neutrophils. We also observe that survival is improved to >50% in the CGD model when mice receive NPs post-infection. This work highlights the application of NPs to improving outcomes to acute bacterial infection in CGD, demonstrating the translational potential of conditionally-immortalized myeloid progenitors as a cellular therapy.

## Introduction

Neutrophils are one of the first responding leukocytes to infection and play a pivotal role in host defense through a wide range of antimicrobial effector functions (1). Chronic granulomatous disease (CGD) is an inherited disorder that is the most common primary immunodeficiency affecting neutrophil function. Those with CGD are highly susceptible to bacterial and fungal infection, especially by *Staphylococcus aureus* (2). Despite such immunocompromise, CGD is also marked by a hyper-inflammatory phenotype that pathologically manifests in the formation of either infectious or sterile granulomas in the lungs, skin, and gastrointestinal tract (3). Current management of CGD includes prophylactic treatment with itraconazole and trimethoprim/sulfamethoxazole to prevent infections and corticosteroid use for granulomatous complications (2). Still, the rise of antibiotic-resistant microbes presents a challenge for preventing and treating infection (4–8). CGD patients who reach adulthood face morbidity due to chronic inflammation, including high rates of pulmonary fibrosis and abnormal pulmonary function (9, 10).

Hematopoietic stem cell transplantation (HSCT) is considered curative for many primary immune diseases, but is currently pursued sparingly because of the infectious risk associated with the required bone marrow conditioning by myeloablation (11–14). The infectious risk of HSCT for CGD patients in particular is concerning, with a recent retrospective multicenter study finding that infection was responsible for 42% of mortality within the first 3 years post-transplant (11). However, the effectiveness of HSCT demonstrates the value of therapeutics targeting engraftment of functional immune cells.

Here, we investigate an experimental neutrophil progenitor-based cellular therapy that is not dependent on bone marrow conditioning and demonstrate its utility in mediating acute host defense in a mouse model of CGD. This therapy is an extension of the previous work by Wang et al. demonstrating that induced ectopic expression of HoxA9 or HoxB8 in murine bone marrow stem cells conditionally immortalizes myeloid progenitors (15). Myeloid progenitors with enforced HoxB8 expression expand exponentially and differentiate into neutrophils after HoxB8 withdrawal *in vitro*. We have previously demonstrated engraftment of HoxB8-conditional neutrophil progenitors (NPs) in the bone marrow of recipient mice, absent irradiation or chemical conditioning (16). Over the course of several days, engrafted NPs differentiate into neutrophils in the bone marrow and egress into the periphery as mature neutrophils (16, 17). Others have demonstrated the potential for HoxB8-conditional progenitors to prolong survival in the context of fungal infection in mice; however, these prior models depend on conditioning for the engraftment of transfused cells (18, 19). In the present study, we demonstrate donor-derived neutrophil (DDN) antimicrobial effector functions *in vivo* and observe that NP transplantation improves outcomes in CGD mice subjected to respiratory infection with *S. aureus*.

## Results

### Donor neutrophils employ anti-bacterial effector functions *in vivo*

We previously demonstrated that, following their transfusion into naïve recipient mice, HoxB8-conditional NPs engraft in the unconditioned hematopoietic niche, differentiate into mature neutrophils, egress to the periphery, and have effector activities equivalent to those of endogenous neutrophils *ex vivo* (16). Here, we probed the *in vivo* antimicrobial function of DDNs by utilizing a bi-fluorescent *S. aureus* killing assay (20). In a first set of experiments, congenic CD45.1 (wild-type) mice were provided a transplant of CD45.2 NPs and subsequently infected with bi-fluorescent *S. aureus* for 12 hours before harvesting the bronchoalveolar lavage (BAL; Figure 1A). BAL samples were stained with anti-Ly6G and anti-CD45.1 or anti-CD45.2 to identify host and donor neutrophils by flow cytometry (Figure 1B).

**Figure 1.**
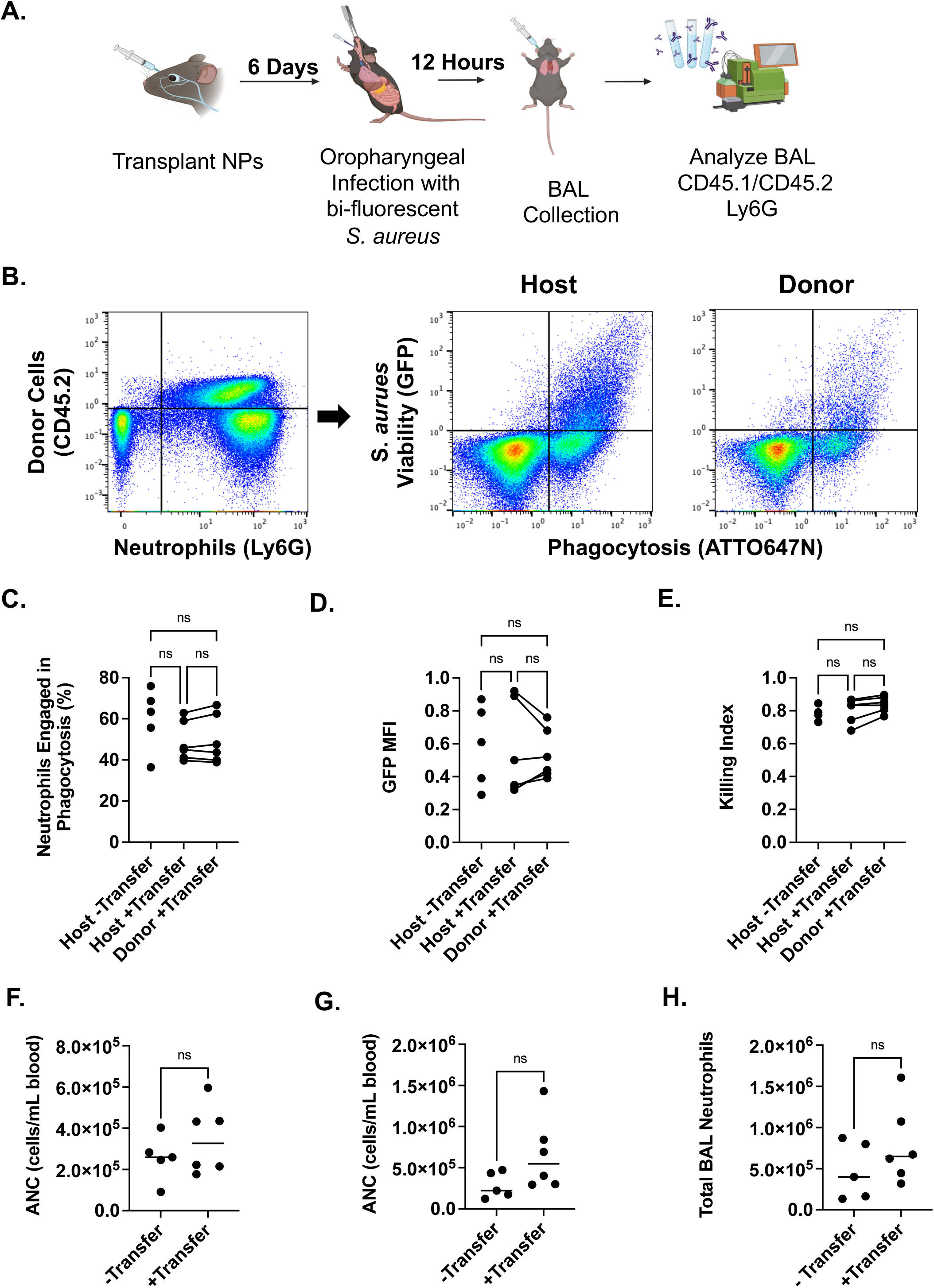
DDN killing capacity of *S. aureus in vivo*. (A) Schematic for evaluating DDN killing of *S. aureus in vivo.* Mice were provided adoptive transfer of NPs 6 days prior to infection with bi-fluorescent *S. aureus*. Twelve hours post infection, mice were sacrificed, and BAL collected for analysis by flow cytometry. Donor cells were identified by expression of CD45.1 vs CD45.2 and neutrophils were identified by Ly6G. (B) Representative flow cytometry analysis of BAL. (C) The percent of neutrophils engaging in phagocytosis was calculated as the percentage of neutrophils that were ATTO647N^+^ 12 hours post-infection. (D) The GFP MFI of host and donor neutrophils represents the intracellular burden of viable *S. aureus* in neutrophils engaging in phagocytosis. (E) The killing index represents the proportion of neutrophils engaging in phagocytosis which have completed killing such that values closer to 1.0 have the highest level of successful killing of *S. aureus*. The absolute neutrophil count (ANC) of CD45.1 mice with or without transplant was measured by flow cytometry in the peripheral blood (F) before and (G) 12 hours after pulmonary infection with *S. aureus*. (H) Likewise, total BAL neutrophil count 12 hours post-infection was measured (n = 5-6, across two independent experiments).

The bi-fluorescent *S. aureus* enables the quantification of phagocytosis and intracellular killing by neutrophils on a single-cell level (20). The two fluorescent markers, ATTO647N and GFP, allow the measurement of neutrophil uptake of *S. aureus* and its viability, respectively. Comparing host and donor neutrophils within BAL of the same mouse, we observed similar engagement in phagocytosis of *S. aureus* between the host and donor neutrophils, which was comparable to neutrophils in the control mouse which did not receive an adoptive transfer of NPs (Figure 1C). There was no significant difference in intracellular bacterial burden of the cells engaging in phagocytosis across groups as measured by the GFP MFI (Figure 1D). The killing index is expressed as the ratio of neutrophils that have completed killing of *S. aureus* relative to the total number of neutrophils that have engaged in phagocytosis. Again, we did not observe a difference in the killing index of host and donor neutrophils (Figure 1E). These results indicate that transplanted NPs generate DDNs with phagocytosis and intracellular killing of *S. aureus in vivo* that was equivalent to that of endogenous wild-type host neutrophils. Comparisons to the neutrophil antimicrobial activity in control mice which did not receive a transplant indicates that the presence of NP-derived neutrophils also did not impact the function of endogenous neutrophils.

To better understand if the transplant of NPs impact endogenous neutrophil dynamics, we also monitored total neutrophil counts in the peripheral blood and BAL. There were no differences in the peripheral absolute neutrophil count (ANC), including donor cells, pre- or post-infection (Figure 1F, G). Likewise, the total count of neutrophils detected in the BAL was comparable between the animals that did or did not receive NP transplantation (Figure 1H). These data suggest that transplantation of NPs does not affect endogenous granulopoiesis and neutrophil recruitment.

### NPs protect CGD mice from pulmonary *S. aureus* infection

Since DDNs can mediate anti-bacterial responses *in vivo*, we reasoned that transplant of NPs may be able to serve as a cellular therapy to support diseases of neutrophil dysfunction. Previous studies of human CGD carriers or chimeras resulting from bone marrow transplantation demonstrate that competent ROS production in as low as 10% of circulating neutrophils can provide sufficient host protection from infection (3, 21). Therefore, we pursued NP transplantation in the *Cybb* knockout murine model of CGD to evaluate the therapeutic potential of NPs (22). Donor cell engraftment kinetics lead to >10% DDN relative to the total population in the periphery by day 6 post-transplant in wild-type (WT) and CGD mice (Supplemental Figure 1A). Subjecting the transplanted mice to a pulmonary *S. aureus* infection, we saw a similar recruitment of DDNs in WT and CGD mice at 24 hours post-infection (Supplemental Figure 1B).

To investigate the ability of DDNs to protect CGD mice, we infected CGD and WT mice 6 days after they received a transplant of NPs. Mice received a pulmonary infection via a bolus of 8×10^7^ CFU *S. aureus*, an inoculum that led to severe infection in CGD mice and mortality within the first 3 days post-infection (Figure 2A). Mice were monitored for humane endpoints until 3 weeks post-infection to assess for potential rebound infections. Control groups included mice that received a saline injection or a transplant of *Cybb*^-/y^ NPs. For clarity, we will refer to the recipient mice that were treated and infected as WT or CGD and the progenitors transplanted as *Cybb^+/y^* NPs or *Cybb^-/y^*NPs.

**Figure 2.**
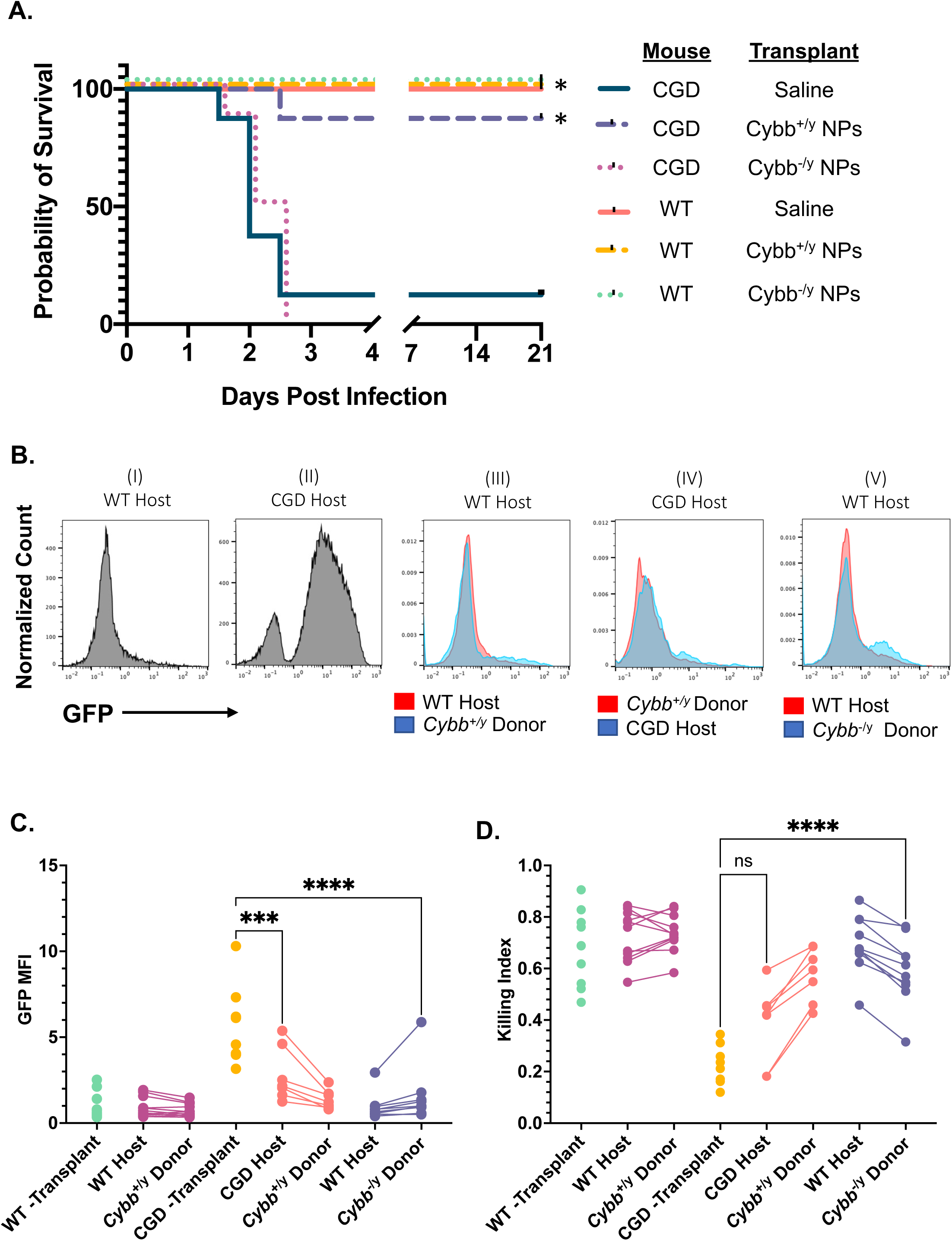
Effect of NP transplant on *S. aureus* pneumonia in the unconditioned murine model of CGD. (A) Mice were treated six days before infection by pulmonary inoculation of *S. aureus* and monitored for survival. While the experiment was terminated after three weeks, all mortality events occurred within the first three days. \**p* < 0.05, comparison to CGD/saline group. For a statistical comparison of all groups see Supplemental Figure 2 (5 independent experiments; n = 5 for WT recipients, n = 8 for CGD recipients). (B-D) Mice, as indicated, were infected with bi-fluorescent *S. aureus* and BAL was collected 12 hours later. (B) Representative flow cytometry histograms of GFP signal of BAL neutrophils (Ly6G^+^), host or donor by CD45.1 or CD45.2, among those engaged in phagocytosis (ATTO647N^+^) in the following groups: (I) CD45.1 non-transplanted mouse, (II) CGD non-transplanted mouse, (III) CD45.1 host with CD45.2 *Cybb^+/y^* NP transplant, (IV) CD45.2 CGD host with CD45.1 *Cybb^+/y^* NP transplant, and (V) CD45.1 host with CD45.2 *Cybb^-/y^* NP transplant. (C) MFI of GFP in ATTO647N^+^ host or donor neutrophils. (D) The proportion of *S. aureus* containing neutrophils that have killed the internalized *S. aureus* twelve hours post-infection (n = 6-11, across 8 independent experiments).

Within the first 3 days post-infection, nearly all untreated CGD mice succumbed to infection (Figure 2A, Supplemental Figure 2). However, the transplant of *Cybb*^+/y^ but not *Cybb*^-/y^ NPs led to a rescue to near WT levels of survival in the CGD mice (Figure 2A, Supplemental Figure 3). As expected, the survival of WT mice with an already competent host defense response was unaffected by the transplant of either *Cybb*^+/y^ or *Cybb*^-/y^ NPs (Figure 2A, Supplemental Figure 2).

### WT DDNs enhance *S. aureus* intracellular killing by CGD neutrophils *in vivo*

To better understand the ability of NP transplantation to improve the survival of CGD mice subjected to pulmonary *S. aureus* infection, we again employed the bi-fluorescent *S. aureus* to assess antimicrobial function at the level of single neutrophils. As expected, at 12 hours post-infection, CGD mice have a large population of neutrophils with viable intracellular *S. aureus*, as visualized by the predominantly GFP^+^ neutrophils among those engaging in phagocytosis (Figure 2B, subpanel II). By contrast, WT mice had predominantly eradicated the internalized *S. aureus* with a dramatic decrease in the GFP fluorescence within phagocytosing neutrophils (Figure 2B, subpanel I). Using the congenic transplant of either *Cybb^+/y^* or *Cybb^-/y^* NPs, we compared the ability of CGD and WT endogenous neutrophils and DDNs to kill *S. aureus* in the chimeric models. Surprisingly, we observed a shift in the capacity of CGD neutrophils to kill *S. aureus* both in CGD mice that received *Cybb^+/y^* donor NPs and in *Cybb*^-/y^ DDNs in CD45.1 WT recipient hosts (Figure 2B, subpanels IV-V). Quantifying *S. aureus* intracellular clearance by GFP MFI, there was an improvement in the antimicrobial function of CGD neutrophils in the mixed chimeric setting compared to the CGD neutrophils in the BAL of animals that did not receive NP transplant (Figure 2C, Supplemental Figure 3). Similarly, there was a significant increase in the killing index in the *Cybb^-/y^* donor neutrophils compared to host CGD neutrophils in the non-transplanted model (Figure 2D, Supplemental Figure 3). While not reaching significance at the group sizing evaluated, the killing index average trended towards an improvement in the host CGD cells in chimeric mice (0.38) compared to the CGD cells of the non-transplanted CGD host (0.22, Figure 2D, Supplemental Figure 3). Despite the increase in the antimicrobial capacity of CGD neutrophils that was observed, when analyzing groups in a pairwise manner, CGD neutrophil function was not completely restored to WT levels (Supplemental Figure 3). However, these data suggest a robust improvement in the antimicrobial activity of CGD neutrophils in the co-presence of WT neutrophils compared to their activity in isolation.

### NPs do not fully rescue neutropenic mice from *S. aureus* pulmonary infection

The findings that WT neutrophils may augment CGD neutrophil clearance of *S. aureus* led us to question whether the therapeutic effect of *Cybb^+/y^*NPs is dependent on the improved function of the normally impaired CGD host neutrophils. We therefore undertook experiments to determine if NPs could rescue severe infection in neutropenic mice.

NP transplant protection from neutropenia can be evaluated through depleting neutrophils by administering an anti-Ly6G antibody to mice (23). Through engineering a Ly6G knockout (*Ly6G^-/-^*) NP cell line using CRISPR, transplanted NPs and their DDNs should be shielded from depletion (23). Loss of Ly6G expression did not influence the maturation of NPs into neutrophils *in vitro* based on the acquisition of CXCR2 and lost expression of cKit (Supplemental Figure 4A, B). *In vivo*, the *Ly6G^-/-^* DDNs (CD45.1^-^ CD45.2^+^) in the periphery had similar expression of neutrophil markers CD11b and CXCR2 as the Ly6G^+^ host neutrophils (CD45.1^+^CD45.2^-^, Supplemental Figure 4C).

To evaluate the impact of *Ly6G^-/-^* NPs on the survival of neutropenic mice subjected to respiratory infection, WT mice were first provided an adoptive transfer of *Ly6G^-/-^* NPs. The mice were then injected with anti-Ly6G at 6 days post-transplant to deplete endogenous neutrophils and then infected with *S. aureus* the next day (Figure 3A). Over the subsequent three weeks survival was monitored for the three study groups: WT mice treated with *Ly6G^-/-^*NPs and subjected to anti-Ly6G depletion (treated group), untreated WT mice subjected to anti-Ly6G depletion (untreated group), and untreated WT mice that were not depleted of their neutrophils (undepleted control group). As expected, there was a significant decrease in the survival of untreated neutropenic mice compared to the undepleted control group (Figure 3A). However, the treated group was not statistically different from either the untreated group or the undepleted control group. Thus, NP transplantation in neutropenic mice may not provide as high a level of functional host defense against *S. aureus* as it does in CGD mice.

**Figure 3.**
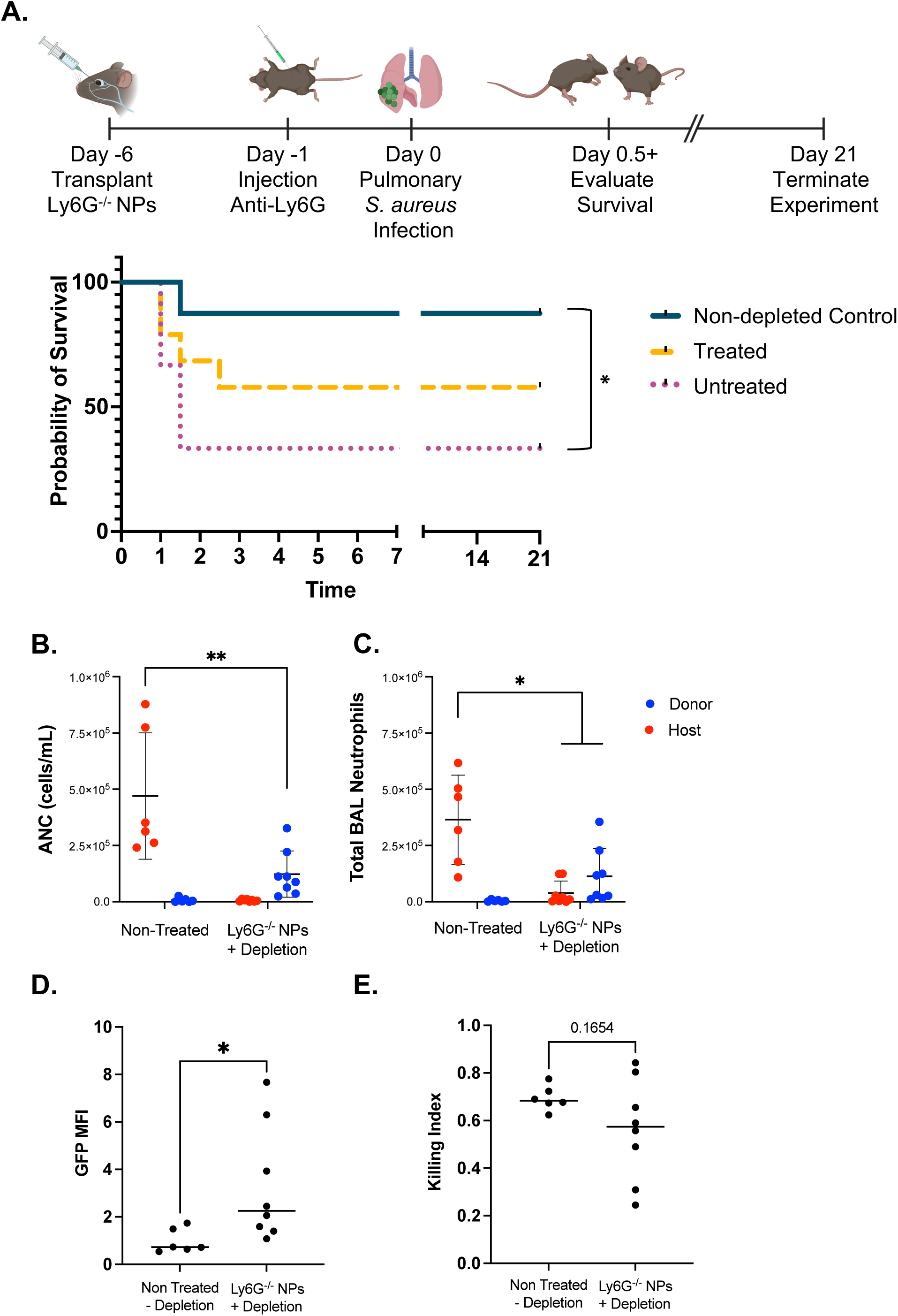
Impact of *Ly6G^-/-^*NP transplant on *S. aureus* pneumonia in anti-Ly6G depletion murine model of neutropenia. (A) Schematic timeline for the neutrophil depletion survival study. C57BL/6 mice were treated by transplantation of *Ly6G^-/-^* NPs 6 days prior to infection with *S. aureus*. To deplete endogenous host neutrophils, mice received anti-Ly6G 24 hours prior to infection. Survival was monitored over the subsequent 3 weeks. There was a statistically significant difference in survival between the non-depleted control group (n = 8) and the untreated group that did not receive NPs (n = 18; Log-rank test, *p* = 0.0134). However, there was no statistical difference observed between the non-depleted control group and the NP-treated group (n = 19; *p* = 0.149), or between the untreated and treated group (*p* = 0.127). (B) Blood and (C) BAL were analyzed by flow cytometry to determine total counts of *Ly6G^-/-^* DDN (CD45.2^+^) and host neutrophils (Ly6G^+^). (D) The bacterial burden (GFP MFI) and (E) proportion of cells completing killing (the killing index) of WT neutrophils in the BAL of naïve mice infected with bi-fluorescent *S. aureus*, compared to the DDN in mice treated with *Ly6G^-/-^* NPs and subjected to anti-Ly6G depletion prior to infection (n = 6-8 per group, across 4 independent experiments).

To gain insight into the function of the *Ly6G^-/-^* DDNs *in vivo*, we employed the bi-fluorescent *S. aureus* model. Untreated CD45.1 mice and mice provided an adoptive transfer of *Ly6G^-/-^*NPs were subjected to anti-Ly6G neutrophil depletion and then inoculated with pulmonary bi-fluorescent *S. aureus* following the same initial timeline as the survival studies. The blood and BAL was collected 12 hours post-infection to evaluate ANC and the killing capacity of the *Ly6G^-/-^* DDNs. The anti-Ly6G depletion resulted in near complete loss of endogenous host neutrophils in the periphery and BAL (Figure 3B,C). However, despite the contribution of *Ly6G^-/-^*DDNs, there were significantly fewer total neutrophils in both the blood and BAL of the mice subjected to depletion (Figure 3B,C). We observed that while the GFP MFI of *Ly6G^-/-^* DDNs was higher compared to neutrophils in the undepleted control group, indicating an increased viable bacterial burden (Figure 3D), the *S. aureus* killing index of BAL neutrophils was comparable between *Ly6G^-/-^* DDNs in depleted mice and endogenous host neutrophils in undepleted mice (Figure 3E). We attribute the impaired eradication of *S. aureus* by *Ly6G^-/-^*DDNs to the decreased total neutrophil count observed in the blood and BAL of the depleted mice (Figure 4B,C). This effectively leads to a higher multiplicity of infection (MOI), which can lead to decreased *S. aureus* clearance by functionally competent neutrophils (24).

**Figure 4.**
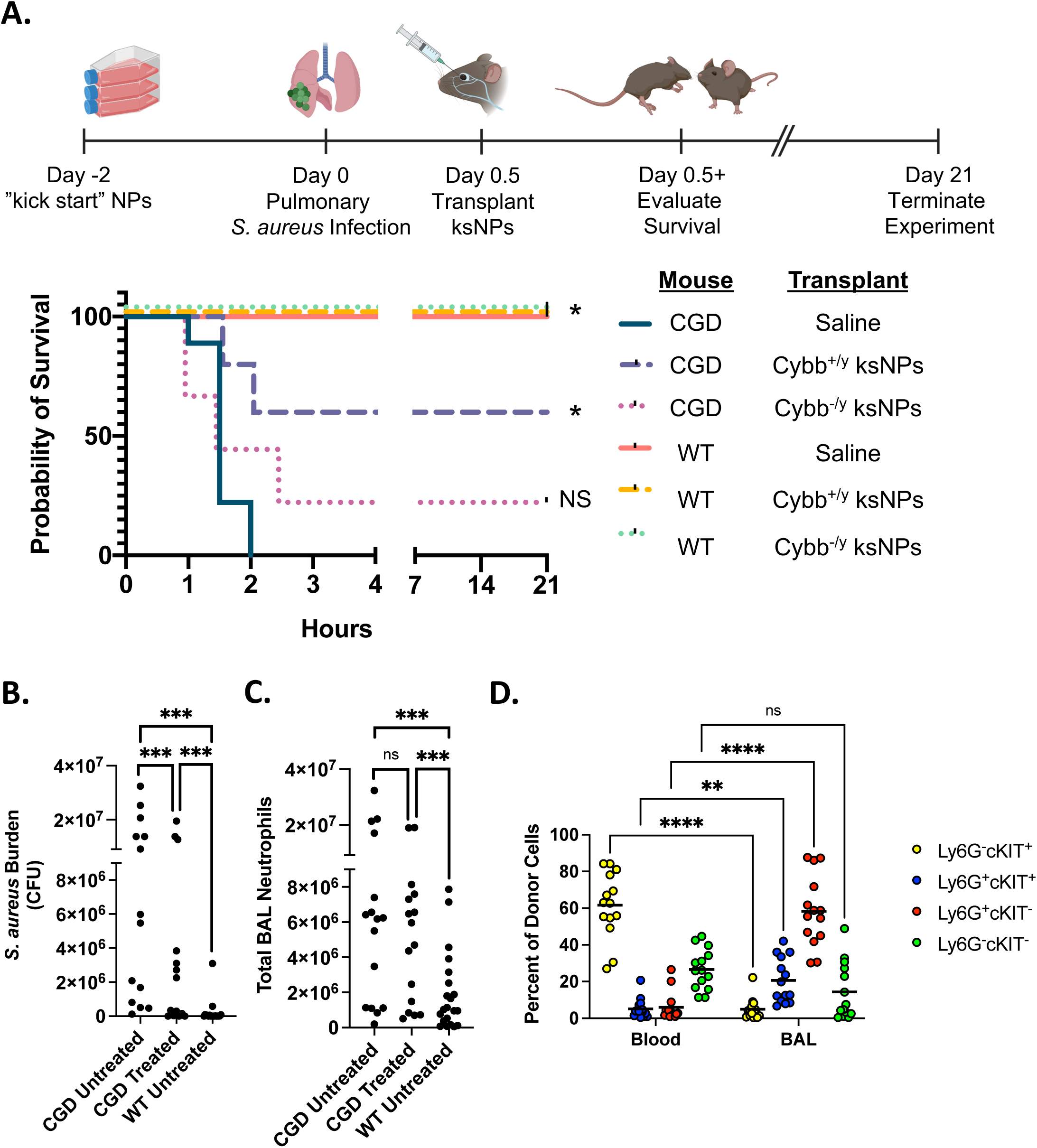
Influence of ksNPs on CGD immune response to *S. aureus* 12 hours post-transplant. (A) Mice were treated with ksNPs 12 hours prior to pulmonary infection with *S. aureus* and then evaluated for survival. Mice were monitored for three weeks post-infection (5 independent experiments). For a statistical comparison of all groups see Supplemental Figure 2. (B) *S. aureus* bacterial burden measured by growth on TSB-agar plates and (C) total neutrophil (Ly6G^+^) counts determined by flow cytometry in the BAL of treated and untreated CGD mice, and untreated WT mice. (D) Expression of cKit and Ly6G on donor cells present in the periphery and the BAL, measured by flow cytometry (For B-D, n = 15-18/group, across 10 independent experiments).

### NP transplantation during established respiratory *S. aureus* infection rescues CGD mice

The kinetics of NP differentiation *in vivo*— not reaching substantial DDN chimerism in the periphery until about 5 days after NP transplantation (16)— suggest that NP transplantation may have limited efficacy as a cellular therapy for acute infections, as we observed mortality within three days for CGD mice infected with *S. aureus* (Supplemental Figure 1A, Figure 2A). Since NPs remain in a precursor state in the presence of the ectopic HoxB8 transcription factor, we hypothesized that providing progenitors a “kickstart” towards initiating the maturation process by removing the HoxB8 inducing agent, tamoxifen, from culture several days prior to NP transplant—as opposed to at the time of NP transplant—might lead to an acceleration of their differentiation *in vivo* and allowing their therapeutic use post-infection. We therefore used this approach to perform experiments using “kickstart” neutrophil progenitors (ksNPs) that were cultured without tamoxifen for 3 days prior to transplantation. We found that when mice were infected and then provided the ksNPs 12 hours later, a robust population of DDNs appeared in the circulation of WT mice by day 4 post-transplant (Supplemental Figure 5A, B). Interestingly, the surface expression level of cKit and lack of Ly6G expression on ksNPs was equivalent to that of NPs in the presence of tamoxifen, suggesting that ksNPs are still in a progenitor state at 3 days after relieving ectopic HoxB8 expression (Supplemental Figure 5C).

We next looked at the ability of treatment with ksNPs post-infection to rescue CGD mice from pulmonary infection with *S. aureus*. In these studies, ksNPs were transplanted 12 hours after WT and CGD mice were infected with *S. aureus* (Figure 4A, Supplemental Figure 2). The treatment and control groups for this experiment were the same as those employed in the prior CGD survival studies (Figure 2). All WT mice, regardless of the treatment, survived infection at this dose of *S. aureus* (Figure 4A, Supplemental Figure 2). There was no significant difference in survival between the mice treated with *Cybb*^-/y^ ksNPs and control CGD mice that received a saline injection instead of ksNPs. However, there was a significant improvement in the survival of CGD animals treated with *Cybb*^+/y^ ksNPs (60%) compared to the control group of CGD mice that did not receive ksNPs (0%, Figure 4A).

### Improved mortality of infected CGD mice is mediated by a small population of DDN’s influence on bacterial burden and inflammation

To further probe the mechanism of improved survival of infected CGD animals treated with ksNPs, we performed analyses of mice at 24 hours post-infection, 12 hours post-transplant. We observed that the bacterial burden in the BAL was significantly lower in CGD mice that received ksNPs versus untreated CGD mice (Figure 4B). The transplant did not appear to affect the hypercellularity associated with CGD infection, as both the treated and untreated CGD mice had significantly more neutrophils in the BAL compared to the WT control (Figure 4C). However, we were surprised to find that in the treated mice, only 0.63% (+/-0.67%) of BAL neutrophils were the *Cybb^+/y^* DDN (Supplemental Figure 6A).

To further understand these observations, we evaluated the characteristics of the ksNPs in the blood and BAL of infected CGD mice. The DDN in the BAL often had a Ly6G^+/med^ and Ly6G^+/high^ population (Supplemental Figure 6B). Surprisingly, we found that the donor-derived cells in the blood were primarily in an immature state (Ly6G^-^cKit^+^) despite the presence of predominantly differentiated DDNs in the BAL (Ly6G^+^cKit^-^; Figure 4D). Interesting there was also a large population Ly6G^+^cKit^+^ cells in the BAL compared to the blood. These data suggest that the immature Ly6G^+^cKit^+^ donor cells may be trafficking to the lung and differentiating locally into mature neutrophils.

Since CGD has hyperinflammatory disease sequelae, we also probed the impact that ksNPs or DDNs may have on the inflammatory response at this time point. We quantified the following cytokines: IL-1⍺, IL-1β, IL-6, IL-10, IL-17⍺, IL-23, G-CSF, IFN-Ɣ, MCP-1, KC, TGF-β1, and TNF-⍺ (Supplemental Figure 7A-S). IL-10 and IL-23 were below the detectable threshold of this assay for most samples and could not be measured, as was the level of IL-1β and TGF-β in the BAL samples. Most cytokines in the serum or BAL were either elevated in both CGD groups, independent of treatment, or indistinguishable across groups. However, there was a significant difference in TNF-⍺ and IL-17⍺ in the serum across all three groups, suggesting that treatment with ksNPs leads to a decrease in these cytokines, but does not restore it to WT levels (Supplemental Figure 7A, C).

## Discussion

Cellular therapies are gaining traction as an appealing approach to improving disease outcomes. The NP cell therapy described here is distinct from current cell therapies and addresses key limitations of the existing neutrophil-based cellular therapies. This NP cell therapy approach involves the adoptive transfer of NPs, which engraft in the bone marrow of unconditioned mice (16). This is a similar process as HSCT; however, unlike HSCT, myeloablation is not required to achieve NP engraftment. This is significant because myeloablation nonspecifically clears the bone marrow of host stem cells, whether the cells are functional or not, leading to a severe state of immunocompromise (25, 26). Alternatively, granulocyte transfusions, like NPs, do not require myeloablation, transiently replenish neutrophils in the periphery, and could be used for similar applications. However, granulocyte transfusions are a controversial approach to treating infection because they have not been shown to improve outcomes in large-scale clinical trials (27). Granulocyte transfusions rely on leukapheresis to collect neutrophils from a healthy donor. The process of collection and storage may have detrimental effects on neutrophil function compromising their ability to clear infections when transplanted (28). Conversely, the supply of NPs is easily maintained through expansion in culture and cryopreservation, eliminating the need for healthy donor participants (15). Also, granulocytes need to be transplanted in high numbers, daily, rendering this therapy impractical for large-scale use (27). Unlike granulocytes, NPs can expand and differentiate *in vivo* supporting the peripheral population for up to a week (16). Given the promising characteristics of NP cell therapy compared to currently available approaches, we propose the further development of HoxB8 NPs as a cellular therapy to treat primary immunodeficiencies, using CGD as a model system.

We found that engraftment of transplanted NPs in unconditioned CGD mice led to a robust population of ROS-producing neutrophils, capable of protection from an otherwise lethal *S. aureus* pulmonary infection. Our analyses of neutrophil intracellular killing of *S. aureus* on a single-cell level suggest that this protection is at least in part due to the ability of the DDNs to confer antimicrobial functional capacity to the CGD neutrophils *in vivo*. This is further validated by the inability of the *Ly6G^-/-^* NPs to rescue experimentally-induced neutropenic mice from a similar infection. Others have demonstrated the transfer of ROS between WT and CGD cells mediated the killing of *Aspergillus fumigatus* (29). Recently, other examples of neutrophil cooperation have been described, including the transfer of substrates between neutrophils deficient in various enzymes within the LTB4 synthesis pathway (30–32). In our study, we speculate that the ability of donor-derived neutrophils to amplify the otherwise deficient host neutrophils’ effector functions may explain why DDNs provide a high level of protection in our CGD model, but were unable to fully rescue our anti-Ly6G neutropenic model, nor the lethal fungal infection in neutropenic mice as investigated by others (19).

Our results demonstrating the efficacy of ksNPs that are transplanted in the context of an ongoing respiratory infection imply the translational potential of this cellular therapeutic approach. For example, ksNP transplant may be a treatment option for drug-resistant infections after they present in the clinic. Since donor cells are present as terminally-differentiated neutrophils in the BAL, but not the peripheral blood, at 12 hours after transplant, it is plausible that they are recruited as progenitors to the lungs and locally undergo maturation. This mechanism of ksNP recruitment to sites of infection and maturation in response to local signals is in alignment with what others have observed. Kim et al. found that cKit^+^ myeloid progenitor cells are recruited from the bone marrow to cutaneous *S. aureus* infections and locally differentiate to generate mature neutrophils (33). They showed that the locally-derived neutrophils have normal effector functions and were critical for animal survival. They attribute the maintenance of the neutrophil population in the abscess to the proliferation and differentiation of the progenitors, and an antiapoptotic cytokine environment (33). For our CGD therapeutic model, follow-up studies are needed to determine if NP engraftment in the bone marrow is a dispensable step and to define the signals that mediate local differentiation of the NPs into mature neutrophils.

Effects of NP transplant on the hyperinflammatory environment observed in CGD warrant further investigation. While we found that NP transplantation results in some significant differences in cytokine levels of TNF-α and IL-17α, overall, the variability in the cytokine concentration in this infection model makes it difficult to draw definitive conclusions or distinguish if there are subtle, but biologically meaningful, differences that are not detected with this sample size. To better characterize potential changes in the cytokine environment with NP treatment, a more chronic infection model may prove superior with a greater degree of resolution between treated and untreated mice.

Applications of NP therapy to immunodeficiencies face a lower barrier to efficacy, compared to infections in otherwise healthy individuals, since many of the infectious burdens are opportunistic and likely could be cleared by uncompromised donor neutrophils. However, applications of cellular therapies may be expanded to the immunocompetent population as well through genetic modifications to NPs that improve the intrinsic function of the neutrophil. The ability to provide this therapy without host conditioning increases the feasibility and practicality of its use in both immunodeficient and uncompromised populations. Our findings provide support for the future of neutrophil-based therapies, particularly in using conditionally-immortalized systems analogous to the murine HoxB8*-*conditional NPs.

## Materials and Methods

### Progenitor cell derivation and culture

Neutrophil progenitors were conditionally immortalized as previously described (15–17). Bone marrow (BM) was aseptically flushed with PBS from a wild type (C57BL/6J), CGD (B6.129S-*Cybb*^tm1Din^/J), or CD45.1 (B6.SJL-*Ptprc*^a^*Pepc*^b^/BoyJ) mouse tibia and femur. Progenitor cells were enriched using an EasySep mouse hematopoietic progenitor cell isolation kit (Stem Cell Technologies). Cells were grown in RPMI (Gibco) supplemented with 10% fetal bovine serum (FBS, Gemini Bio-Products), 1x penicillin/streptomycin (Gibco), 1x non-essential amino acids (Gibco), with 20 ng/mL each of recombinant murine SCF, IL-3 and IL-6 (BioLegend) for three days. Cells were then spin-infected with HoxB8/hygro lentivirus derived from the supernatant of transfected Lenti-X HEK293T cells (Takara Bio). After 24 hours, cells were reseeded in the progenitor media described below with 200 μg/mL hygromycin (Tocris Bioscience) until the uninfected control was no longer viable (about two weeks).

Progenitor cells were maintained in Opti-MEM I reduced serum medium supplemented with: GlutaMax (Gibco), 30 mM beta-mercaptoethanol (Sigma-Aldrich), 10% FBS (Gemini Bio-Products), 1x penicillin/streptomycin (Gibco), 1x non-essential amino acids (Gibco), 50 ng/mL SCF, and 100 nM Z-4-hydroxytamoxifen (Tocris Bioscience). The SCF was derived in-house from a recombinant murine SCF-producing CHO cell line, kindly provided by Dr. Patrice Dubreuil (Centre de Recherche en Cancérologie de Marseille). The ksNPs were prepared by washing NP cells extensively with PBS and resuspending them in progenitor media devoid of Z-4-hydroxytamoxifen. To differentiate progenitors for *in vitro* studies, NPs were washed 3x with PBS and seeded in media devoid of Z-4-hydroxytamoxifen and containing 20 ng/mL SCF and G-CSF. After 2-3 days, cells were re-seeded in media containing 20 ng/mL G-CSF (without Z-4-hydroxytamoxifen or SCF). Differentiated cells were used in assays on days 5-6.

To generate the Ly6G-deficient NP cell line, we used the pLentiCRISPR v2 vector, a gift from Feng Zhang (Addgene plasmid #52961) with the following 20-nucleotide single-guide RNA targeting sequence inserted: CACACAGTAGGACCACAAGA (*Ly6g*). To express Cas9 and sgRNA, NPs were transduced via lentivirus, as described above.

### Animal care and breeding

All studies were performed under the approval of the Lifespan Animal Welfare Committee (Office of Laboratory Animal Welfare Assurance #A3922-01). These studies follow Public Health Service guidelines for animal care and use. CGD mice were first acquired from Jackson Laboratories (Bar Harbor, ME), strain B6.129S-*Cybb*^tm1Din^/J, and used for in-house breeding. This strain was originally developed by disruption of the *Cybb* gene (gp91^phox^) to recapitulate the CGD phenotype (22). Female mice heterozygous for the mutated *Cybb* gene (x-linked) were bred with wild-type males to yield hemizygous wild-type (*Cybb*^+/y^) or knockout (*Cybb*^-/y^) male progeny used in this study. Thus, in these studies employing the mouse model of x-linked CGD, sex as a biological variable was not evaluated. Animals were housed in sterile caging until infection was performed in mice 6-10 weeks of age and provided access to water and standard chow ad libitum.

### NP Transplantation

NPs were pelleted and washed 3x with PBS before being brought to a density of 8 x 10^7^ cells/200 µL normal saline. For the transplant of WT CD45.2^+^ cells into CGD mice (CD45.2 background), donor cells were labeled with Tag-it Violet Proliferation and Cell Tracking Dye (BioLegend) to distinguish donor from host cells. For other groups, the congenic markers CD45.1 and CD45.2 were used to distinguish donor from host cells, as indicated. Mice were anesthetized with isoflurane and 200 µL of NPs or normal saline control was administered via retro-orbital injection.

### Staphylococcus aureus culture and staining

USA300-GFP and USA300-Luciferase strains were a generous gift from Alexander Horswill (University of Colorado). The super folded GFP strain was used for the bi-fluorescent *S. aureus* infection experiments; remaining survival and mechanistic studies were conducted using the non-GFP expressing strain of *S. aureus*. To culture, *S. aureus* from a glycerol stock was grown in tryptic soy broth (TSB, Sigma Aldrich) overnight, shaking at 250 RPM, 37°C. Prior to infection, *S. aureus* was diluted to an optical density (OD_600_) of 0.4 and grown to an OD_600_ of 0.8-1, as measured with a SmartSpec 3000 (Bio-Rad). For bi-fluorescent assays, the USA300-GFP *S. aureus* was stained as previously described (20). Briefly, *S. aureus* at a concentration of 4×10^8^ CFU/mL (OD_600_=1) was stained with 0.5 mg/mL of ATTO647N-NHS ester (Sigma Aldrich) for 30 minutes at room temperature in the dark, then washed 2x with PBS prior to use.

For mouse infection studies, *S. aureus* was washed 2x with PBS and brought to an OD_600_ of 0.5. Cells were pelleted at 2300 x *g* for 10 minutes and resuspended at a ratio of 100 µL/1 mL (concentrated 10x) in normal saline. Animals were anesthetized with isoflurane and a 40 µL bolus (8×10^7^ CFU) of the prepared *S. aureus* was delivered by oropharyngeal aspiration.

### Tissue collection

For analysis of blood, up to 50 µL of blood was collected in an EDTA-coated tube via puncture of the saphenous vein. During terminal procedures, blood was collected via cardiac puncture into an EDTA-coated tube. Prior to antibody staining and analysis by flow cytometry, red blood cells (RBC) were lysed with RBC lysis buffer (BioLegend). Serum was separated by centrifugation for 20 minutes at 2000 x *g*. Bronchoalveolar lavage (BAL) was collected 12 hours post-infection by pushing and withdrawing 3 x 1 mL of sterile PBS/BSA/EDTA (for cytokine studies) or PBS/2% FBS through a catheter inserted into the trachea. Bone marrow was collected by flushing the femur with 1 mL of PBS/2% FBS.

### Flow cytometry analysis

To stain samples for analysis by flow cytometry, cells were suspended in cold FACS buffer (PBS, 2%FBS). Samples were stained for 30 minutes on ice and washed with FACS buffer 2x before acquisition by a MACSQuant Analyzer 10 flow cytometer (Miltenyi). Analysis of results was performed using FlowJo Software (Becton Dickinson). For determination of percent’s, counts, and characterization of host or donor cells, BAL or lysed whole blood was stained with AF647 anti-Ly6G and PE anti-cKit. For staining with the PE anti-Cybb antibody (Santa Cruz Biotechnology, Inc.), cells were first fixed with 4% formaldehyde overnight at 4°C, permeabilized with ice-cold 90% methanol on ice for 1 hour, then washed and stained with anti-Cybb for 1 hour at room temperature.

For the *in vivo S. aureus* killing assay, the BAL from infected mice was stained with anti-Ly6G and anti-CD45.1 or anti-CD45.2. These two antibodies were labeled with the PE or APC-Cy7, results followed the same trends regardless of whether the Ly6G versus CD45.1/CD45.2 antibody was in the PE or APC-Cy7 channel. Each BAL was stained as two separate samples, once with each CD45 antibody. This ensured analysis of host or donor cells was always on a population of cells with the same fluorescent labels.

### Neutrophil depletion

To deplete endogenous neutrophils, 100 µg of purified anti-Ly6G clone 1A8 (Absolute Antibody) was administered to mice via intraperitoneal injection, performed 1 day prior to infection. Control mice received injection of rat IgG2A isotype antibody.

### Colony Forming Unit (CFU) assays

BAL collected from animals 12 hours post-treatment was analyzed for *S. aureus* burden. Plating 3x 10 µL serial dilutions of the BAL on TSB-agar and incubating the plates overnight at 37°C enabled quantification of the total CFU burden. Prior to plating, the BAL was diluted 1:100 in H_2_O pH=11 for 10 minutes to lyse leukocytes. This prevents clumping of the intracellular *S. aureus* on the TSB-agar leading to artificially lower counts (34).

### Cytokine analysis

Cytokine analysis was completed using the LEGENDplex multiplex assay (BioLegend). Experiments were conducted per the manufacturer’s instruction. Analyzed cytokines included: IL-1⍺, IL-1β, IL-6, IL-10, IL-17⍺, IL-23, G-CSF, IFN-Ɣ, MCP-1, KC, TGF-β1, TNF-⍺.

### Statistical analysis

Graphical depictions of the data represent group means and standard deviations (SD) generated using Prism 9 (GraphPad). With the exception of the CFU and cytokine data, statistical analysis was also performed in GraphPad. Statistical significance was determined by ordinary one-way ANOVA to compare groups with an alpha set to 0.05. Alternatively, an unpaired Student’s t-test was utilized when there were only two groups being compared. For data analyzed and reported in Figure 4, a paired Student’s t-test was used when comparing the host and donor cells from the same animal. Indicators of statistical significance are as follows: “ns” for p>0.05, * for p<0.05, ** for p<0.01, *** for p<0.001, and **** for p<0.0001.

Statistical testing for survival curves was completed in GraphPad by calculating the p-value between each group using the Mantel-Cox test and Gehan-Breslow-Wilcoxon test. The calculated p-value was then compared to the adjusted Bonferroni p-value (=(alpha)/(number of comparisons)) with an alpha set to 0.05.

Some statistical analyses were carried out using SAS version 9.4 (The SAS Institute). For CFU and neutrophil count data, generalized linear models for negative binomial data were used to compare conditions. A main effect (covariate) was included for the batch to adjust for day-to-day variation since each batch had representation for all experimental conditions. Classical sandwich estimation was used to adjust for any model misspecification, and Holm test was used to maintain alpha at 0.05 across all group comparisons. For cytokines, it was sufficiently common to have out-of-range readings that generalized Tobit models for lognormal data were used. These also included a covariate for batch to reduce the influence of day-to-day variability and used the Holm test to maintain alpha at 0.05 across all group comparisons. Experiments where the cytokine concentration was outside of the range of detection for all animals were excluded from analysis.

## Author Contribution Statements

K.D.H.: Designing research studies, conducting experiments, analyzing data, and writing the manuscript.

J. T. M.: Statistical analysis of CFU, neutrophil count, and cytokine data.

C.T.L.: Supervising the project, designing research studies, analyzing data, editing the manuscript.

## Acknowledgements

We would like to thank those that provided resources used in these studies, including Patrice Dubreuil (SCF-producing CHO cells), Paul Ekert (inducible HoxB8 plasmid) and Alexander Horswill (strains of USA300). Schematic diagrams were created using BioRender.

