## Supplemental Information for "Neutrophil progenitor cell therapy rescues host defense against *Staphylococcus aureus* in murine chronic granulomatous disease"

All materials and methods are reported in the main manuscript.

**Supplemental Figure 1: NPs contribute to the neutrophil population post-adoptive transfer in the unconditioned CGD murine model.**

NPs were labeled with a fluorescent cell tracker prior to transplant allowing for donor cell identification in the blood and BAL by flow cytometry. The total neutrophil population was identified using anti-Ly6G. (A) The percent of DDNs, relative to total neutrophil population by Ly6G expression, in the peripheral blood of WT and CGD hosts within the first 7 days post-transplant of NPs on day zero. (B) Percent of DDNs in the BAL of WT and CGD animals infected with *S. aureus* on day 7 post-transplant (n = 3 per group, across 2 independent experiments).

**Supplemental Figure 2: Complete Statistical Comparisons of Prophylactic Survival Study (Figure 2A) and Treatment Survival Study (Figure 4A).**

**Supplemental Figure 3: Descriptive statistics of the GFP MFI or killing index results and complete results comparing GFP MFI and killing index across groups.**

**Supplemental Figure 4: *In vitro* analysis of *Ly6G$^{-/-}$* NP differentiation**.

(A) *In vitro* differentiation of WT and *Ly6G$^{-/-}$* NP cells into to neutrophils was marked by increasing CXCR2 expression and decreasing cKIT expression by flow cytometry. WT cells also expressed increased levels of Ly6G which was not detected in the *Ly6G$^{-/-}$* differentiated cells. (B) Percent of cultured cells positive for Ly6G, CXCR2, and cKIT on day 5 of differentiation (n = 3-4, across two experiments). (C) *In vivo Ly6G$^{-/-}$* DDNs express CXCR2 and CD11b at comparable levels to host neutrophils in the peripheral blood, a population of cells that is entirely Ly6G$^{+}$ in the host neutrophils despite lost express in the DDNs (n = 2).

**Supplemental Figure 5: Modulation of Transplant Kinetics.**

(A) Multiple conditions were tested (+/- infection, + NPs or ksNPs) to evaluate time from the transplant to substantial DDN presence in the periphery. Mice provided a transplant in the post-infection group were transplanted 12 hours after pulmonary infection with *S. aureus*. Transplanted mice were harvested for blood 4 days later (n = 5-8, across three independent experiments). (B) Expression of cKit and Ly6G in progenitors with or without the “kick start” towards differentiation (n = 2).

**Supplemental Figure 6: Donor Cell Characterization.**

(A) Percent donor cell population relative to the total neutrophil population in the BAL of mice receiving ksNPs 12 hours post-infection, analyzed at twelve hours post-transplant. (B) Representative flow cytometry plots characterizing the donor cell expression of Ly6G and cKIT in the blood and BAL.

**Supplemental Figure 7: Cytokine Profile in Serum and BAL.**

Cytokine measurements of: (A) TNF-α in the BAL (B) TNF-α in the serum (C) IL-17α in the BAL (D) IL-17α in the serum (E) IL-1α in the BAL (F) IL-1α in the serum (G) IL-6 in the BAL (H) IL-6α in the serum (I) G-CSF in the BAL (J) G-CSF in the serum (K) KC in the BAL (L) KC in the serum (M) MCP-1 in the BAL (N) MCP-1 in the serum (O) IFN-ɣ in the BAL (P) IFN-ɣ in the serum (Q) IL-1β in the BAL (R) TGF-β in the serum. (S) Pairwise statistical comparisons of cytokine data, shown as *p* values.

Supplemental Figure 1

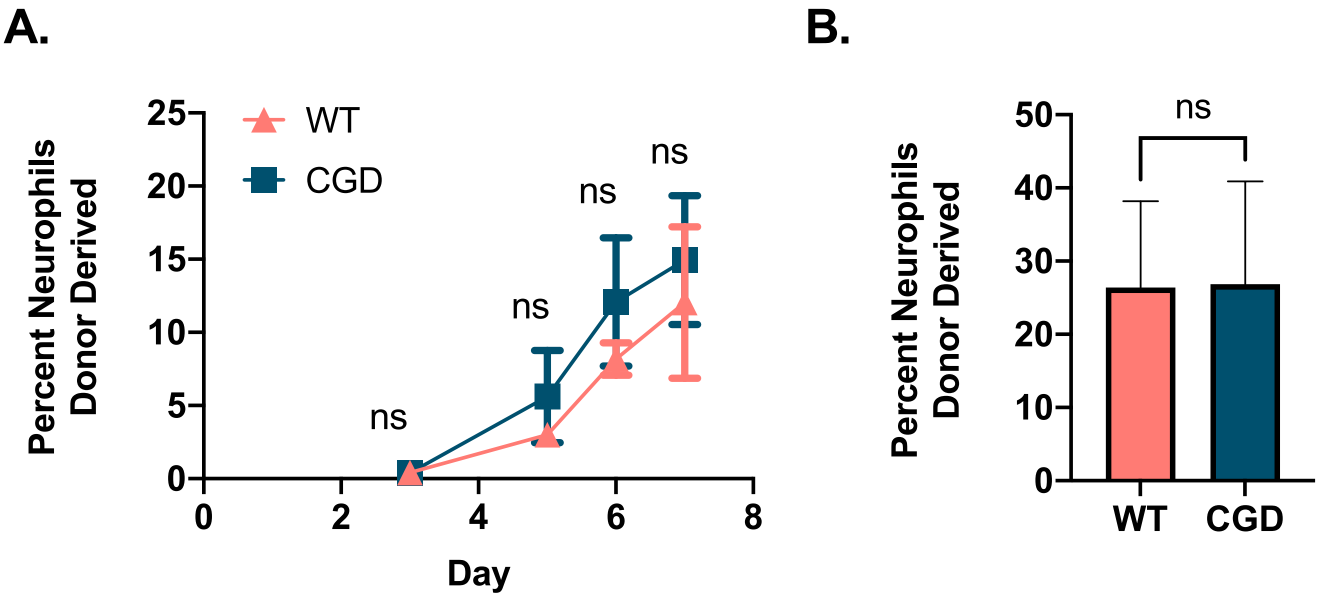

### Supplemental Figure 2

Test Parameters

| | |
|---|---|
| alpha | 0.05 |
| groups | 6 |
| alpha (bonferroni) | 0.0083 |

Prophylactic Treatment (Figure 2)

| Comparison | | Log-rank test | Gehan-Breslow-Wilcoxon | significant? |
|---|---|---|---|---|
| CGD + Saline | CGD + Cybb$^{+/y}$ NPs | 0.0019 | 0.0021 | yes |
| CGD + Saline | CGD + Cybb$^{-/y}$ NPs | 0.9557 | 0.866 | no |
| CGD + Saline | WT + Saline | 0.0044 | 0.0068 | yes |
| CGD + Saline | WT + Cybb$^{+/y}$ NPs | 0.0044 | 0.0068 | yes |
| CGD + Saline | WT + Cybb$^{-/y}$ NPs | 0.0216 | 0.0307 | no |
| CGD + Cybb$^{+/y}$ NPs | CGD + Cybb$^{-/y}$ NPs | 0.0005 | 0.0007 | yes |
| CGD + Cybb$^{+/y}$ NPs | WT + Saline | 0.4292 | 0.4292 | no |
| CGD + Cybb$^{+/y}$ NPs | WT + Cybb$^{+/y}$ NPs | 0.4292 | 0.4292 | no |
| CGD + Cybb$^{+/y}$ NPs | WT + Cybb$^{-/y}$ NPs | 0.5403 | 0.5403 | no |
| CGD + Cybb$^{-/y}$ NPs | WT + Saline | 0.0014 | 0.0026 | yes |
| CGD + Cybb$^{-/y}$ NPs | WT + Cybb$^{+/y}$ NPs | 0.0014 | 0.0026 | yes |
| CGD + Cybb$^{-/y}$ NPs | WT + Cybb$^{-/y}$ NPs | 0.0078 | 0.0148 | no |

ksNP Treatment Statistics (Figure 5)

| Comparison | | Log-rank test | Gehan-Breslow-Wilcoxon | significant? |
|---|---|---|---|---|
| CGD + Saline | CGD + Cybb$^{+/y}$ NPs | 0.0021 | 0.0034 | yes |
| CGD + Saline | CGD + Cybb$^{-/y}$ NPs | 0.2222 | 0.6399 | no |
| CGD + Saline | WT + Saline | <0.0001 | <0.0001 | yes |
| CGD + Saline | WT + Cybb$^{+/y}$ NPs | 0.0002 | 0.0007 | yes |
| CGD + Saline | WT + Cybb$^{-/y}$ NPs | <0.0001 | 0.0003 | yes |
| CGD + Cybb$^{+/y}$ NPs | CGD + Cybb$^{-/y}$ NPs | 0.0813 | 0.0688 | no |
| CGD + Cybb$^{+/y}$ NPs | WT + Saline | 0.0293 | 0.0301 | no |
| CGD + Cybb$^{+/y}$ NPs | WT + Cybb$^{+/y}$ NPs | 0.0884 | 0.0902 | no |
| CGD + Cybb$^{+/y}$ NPs | WT + Cybb$^{-/y}$ NPs | 0.0665 | 0.0679 | no |
| CGD + Cybb$^{-/y}$ NPs | WT + Saline | 0.0005 | 0.0007 | yes |
| CGD + Cybb$^{-/y}$ NPs | WT + Cybb$^{+/y}$ NPs | 0.0055 | 0.0075 | yes |
| CGD + Cybb$^{-/y}$ NPs | WT + Cybb$^{-/y}$ NPs | 0.0029 | 0.0041 | yes |

### Supplemental Figure 3

| | GFP MFI | | Killing Index | |
|---|---|---|---|---|
| | Mean | Stdev | Mean | Stdev |
| WT -Transplant | 1.04 | 0.794 | 0.68 | 0.15 |
| WT Host | 0.901 | 0.747 | 0.69 | 0.11 |
| WT Host 2 | 0.92 | 0.579 | 0.72 | 0.10 |
| *Cybb*$^{+/y}$ Donor 2 | 0.759 | 0.38 | 0.74 | 0.07 |
| *Cybb*$^{+/y}$ Donor | 1.41 | 0.604 | 0.57 | 0.11 |
| CGD -Transplant | 5.65 | 2.5 | 0.22 | 0.08 |
| *Cybb*$^{-/y}$ Donor | 1.48 | 1.6 | 0.59 | 0.13 |
| CGD Host | 2.9 | 1.69 | 0.38 | 0.16 |

| Comparison | Paired | GFP MFI | Killing Index |
|---|---|---|---|
| CGD -Transplant vs. WT Host | No | **** | **** |
| CGD -Transplant vs. *Cybb*$^{-/y}$ Donor | No | **** | **** |
| CGD –Transplant vs. WT Host | No | **** | **** |
| CGD -Transplant vs. *Cybb*$^{+/y}$ Donor | No | **** | **** |
| CGD -Transplant vs. CGD Host | No | ** | ns |
| CGD -Transplant vs. *Cybb*$^{+/y}$ Donor | No | **** | **** |
| *Cybb*$^{-/y}$ Donor vs. CGD Host | No | * | * |
| WT Host vs. CGD Host | No | * | **** |
| *Cybb*$^{+/y}$ Donor vs. CGD Host | No | * | **** |
| *Cybb*$^{+/y}$ Donor vs. WT Host | Yes | ns | ns |
| *Cybb*$^{-/y}$ Donor vs. WT Host | Yes | ns | **** |
| *Cybb*$^{+/y}$ Donor vs. CGD Host | Yes | * | ** |

### Supplemental Figure 4

**A.**

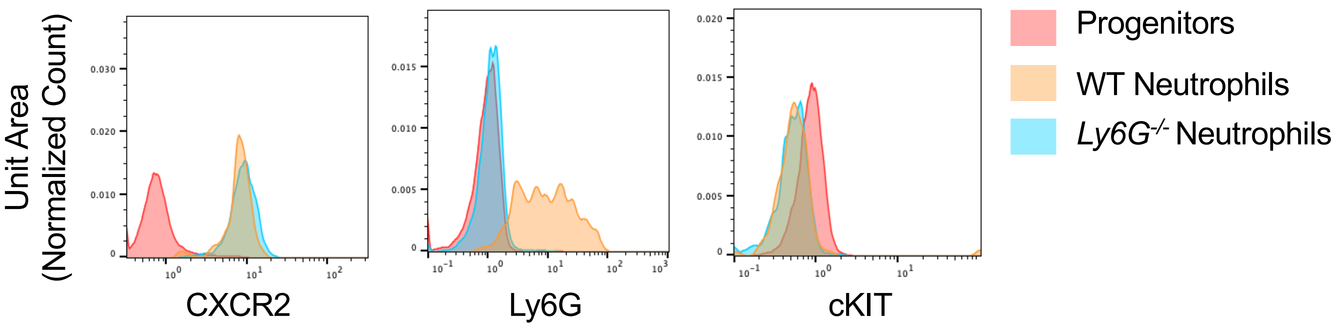


**B.**

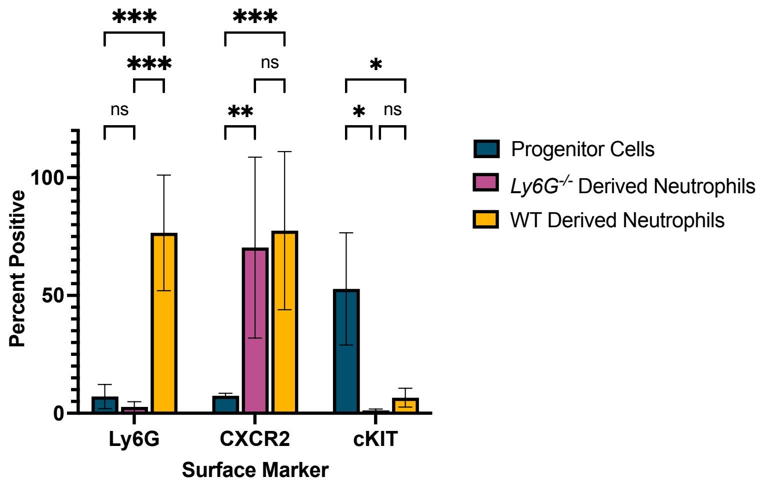


**C.**

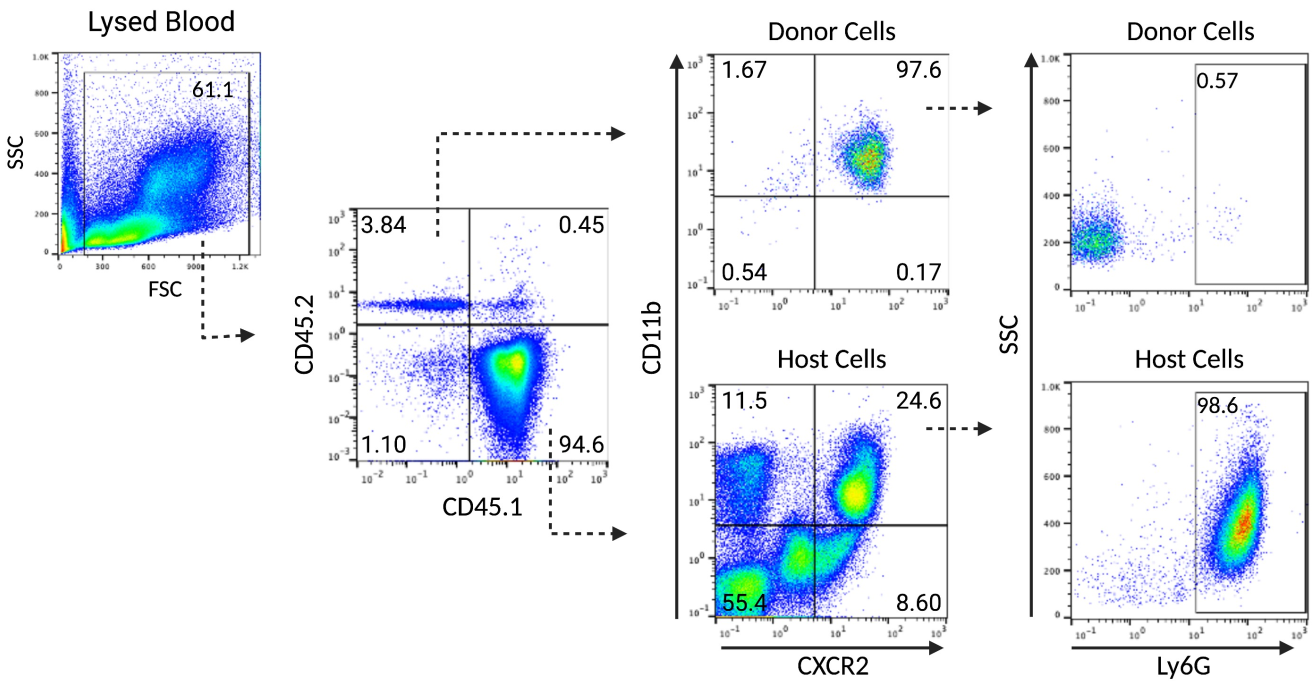

### Supplemental Figure 5

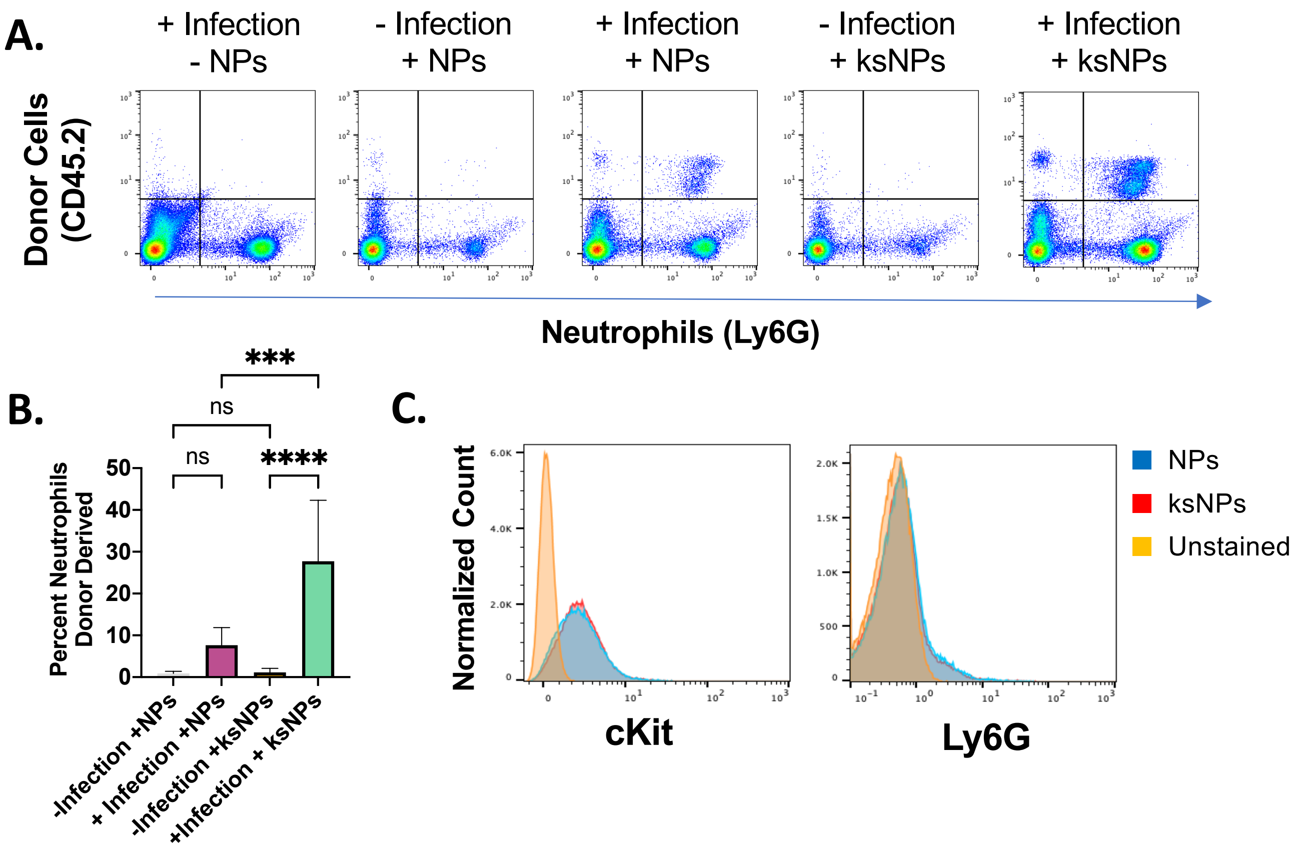

Supplemental Figure 6

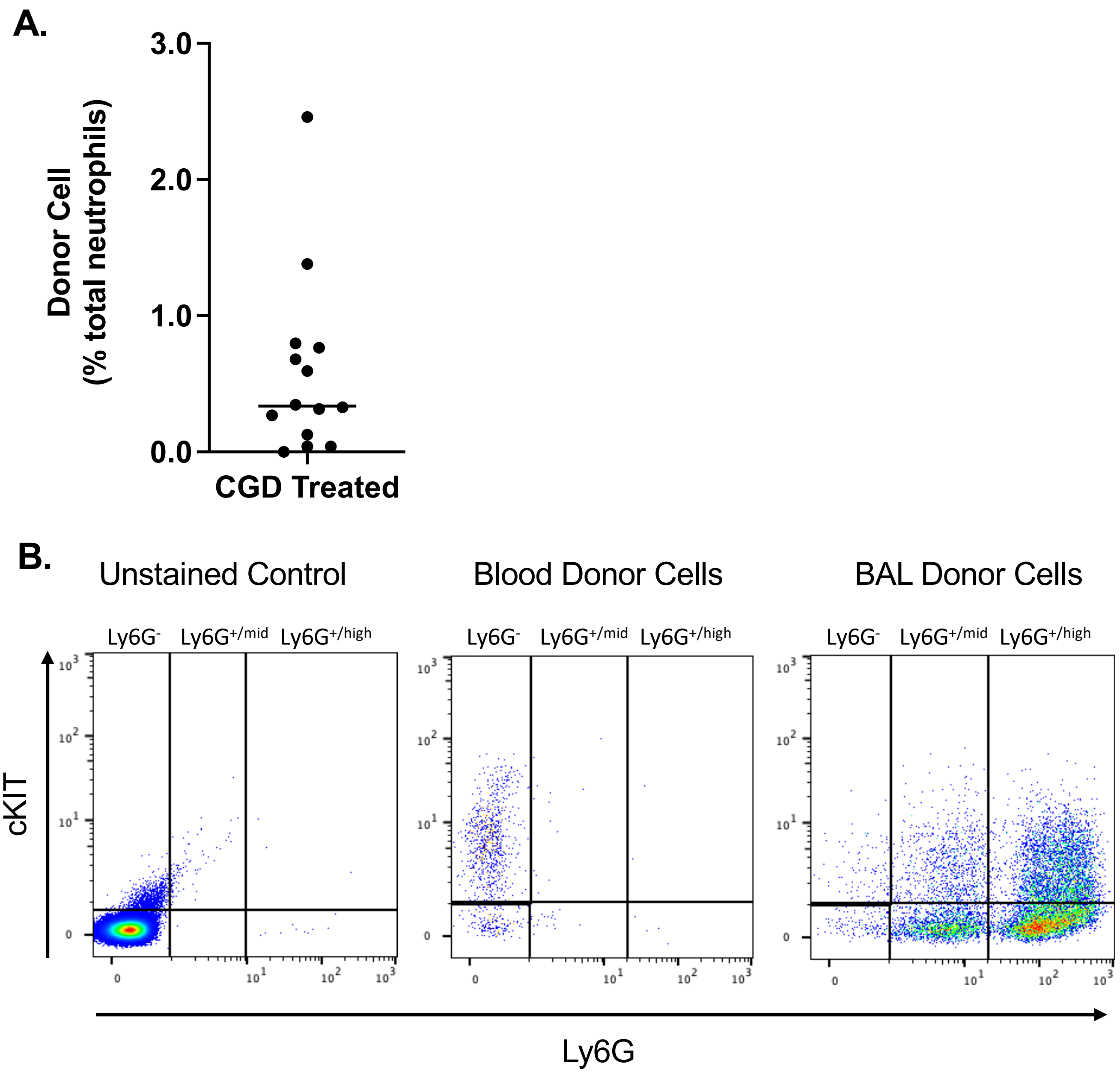

### Supplemental Figure 7

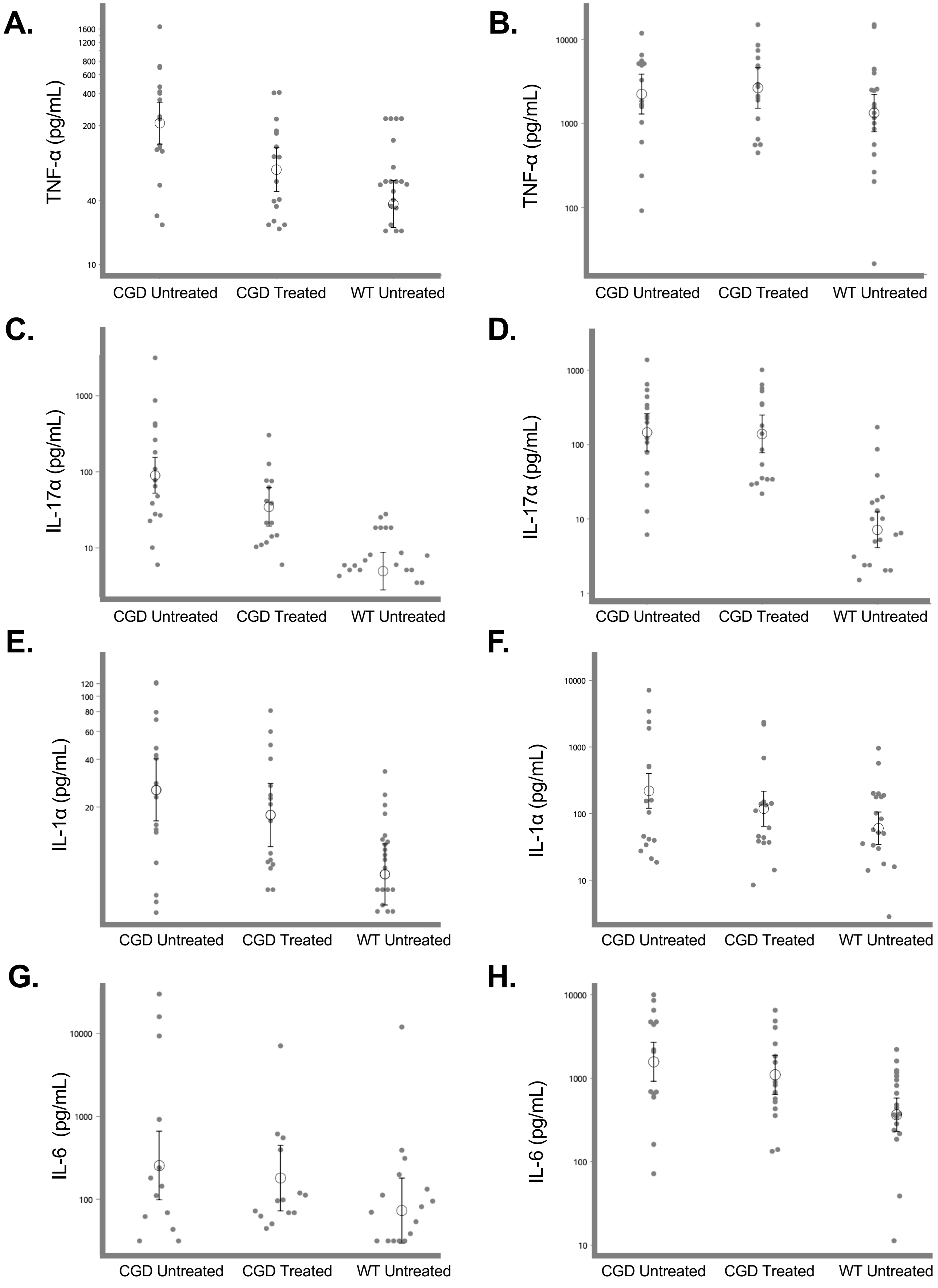

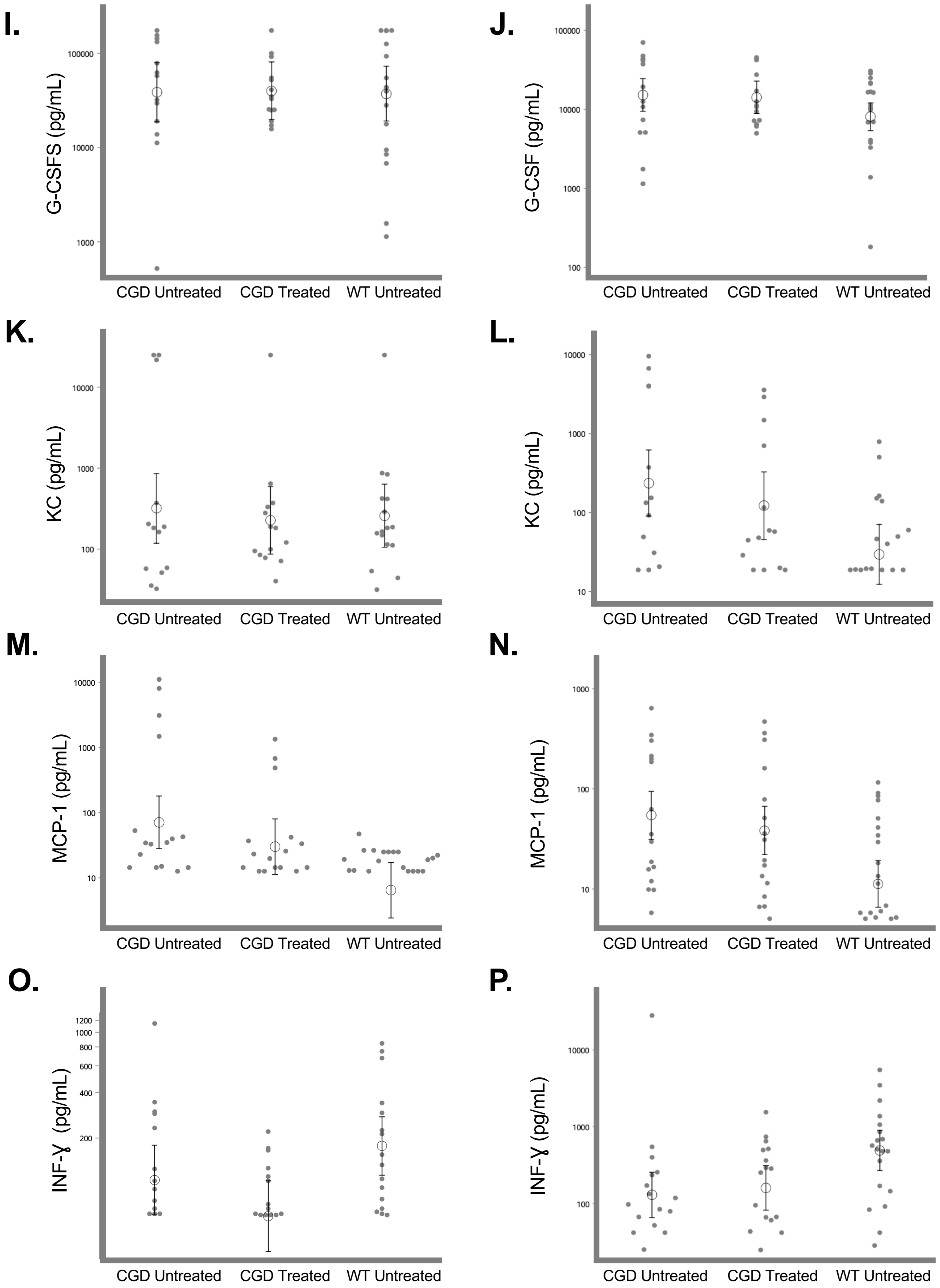
I.
G-CSFS (pg/mL)
100000
10000
1000
J.
G-CSF (pg/mL)
100000
10000
1000
100
K.
KC (pg/mL)
10000
1000
100
L.
KC (pg/mL)
10000
1000
100
10
M.
MCP-1 (pg/mL)
10000
1000
100
10
N.
MCP-1 (pg/mL)
1000
100
10
O.
INF-ɣ (pg/mL)
1200
1000
800
600
400
200
P.
INF-ɣ (pg/mL)
10000
1000
100
CGD Untreated
CGD Treated
WT Untreated

**Q.**

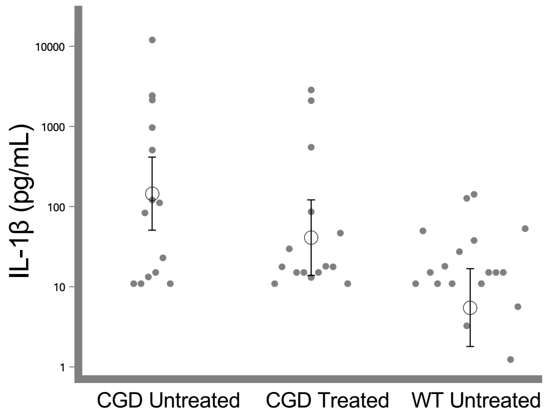


**R.**

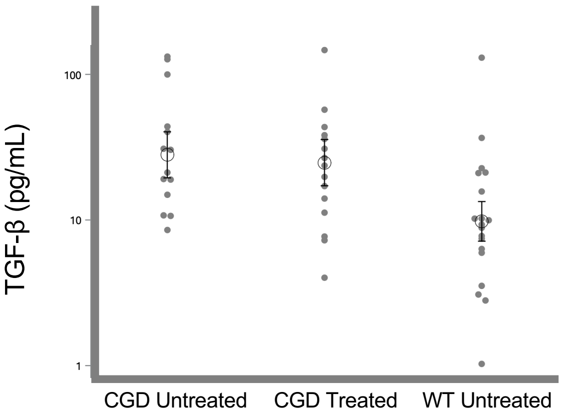


**S.**

| | | CGD TREATED VS CGD UNTREATED | CGD TREATED VS WT | CGD UNTREATED VS WT |
|---|---|---|---|---|
| IL-1α | SERUM | 0.2653 | 0.0149 | 0.0005 |
| | BAL | 0.2108 | 0.2108 | 0.0022 |
| INF-Ɣ | SERUM | 0.2719 | 0.0068 | 0.2719 |
| | BAL | 0.664 | 0.0271 | 0.0127 |
| TNF-α | SERUM | 0.0053 | 0.0305 | <0.0001 |
| | BAL | 0.6724 | 0.2162 | 0.3536 |
| MCP-1 | SERUM | 0.2106 | 0.262 | 0.0004 |
| | BAL | 0.3846 | 0.0034 | 0.0002 |
| IL-17α | SERUM | 0.018 | <0.0001 | <0.0001 |
| | BAL | 0.9172 | <0.0001 | <0.0001 |
| KC | SERUM | 1 | 1 | 1 |
| | BAL | 0.3403 | 0.0711 | 0.0053 |
| IL-6 | SERUM | 0.6019 | 0.3212 | 0.177 |
| | BAL | 0.3479 | 0.005 | 0.0002 |
| G-CSF | SERUM | 1 | 1 | 1 |
| | BAL | 0.8436 | 0.1512 | 0.1411 |
| IL-1β | BAL | 0.1029 | 0.021 | 0.0001 |
| TGFβ | BAL | 0.6365 | 0.0003 | <0.0001 |
